# Spatial Polarization-Induced Fluorescence Fluctuation Imaging (SPIFFI) Enables Single-shot Super-Resolution and Multidimensional Imaging

**DOI:** 10.64898/2025.12.12.693764

**Authors:** Wei Guo, Lely Feletti, Aleksandra Radenovic

## Abstract

Fluorescence super-resolution microscopy has advanced optical imaging into the nanoscale regime, transforming biological and interdisciplinary research. However, conventional wide-field super-resolution techniques often compromise temporal resolution, thereby limiting the ability to capture rapid and transient biological events in living systems. Here, we introduce spatial polarization-induced fluorescence fluctuation imaging (SPIFFI), a multi-channel polarimetric method that enables single-shot super-resolution imaging and six-dimensional information extraction. By leveraging the inherently smaller point spread function (PSF) under polarized detection and capturing polarization-dependent spatial fluctuations across multiplexed channels, SPIFFI achieves instant resolution enhancement from a single exposure. This capability substantially enhances the feasibility of volumetric live-cell super-resolution imaging. Moreover, SPIFFI images can be seamlessly integrated with existing fluctuation-based methods for further post-processing and resolution improvement. We demonstrate the versatility of SPIFFI through experiments on both fixed and live cells, capturing rapid subcellular dynamics and enabling high-throughput, multidimensional imaging beyond the diffraction limit. SPIFFI thus offers a practical and robust platform for real-time super-resolution imaging in biological research.

## 1. Introduction

Super-resolution fluorescence microscopy has revolutionized biological imaging by surpassing the diffraction limit^1-6^, achieving nanometer-scale resolution, and now approaches the sub-nanometer level^7-9^. Although it has yet to reach the structural detail of cryo-electron microscopy, optical super-resolution uniquely enables visualization of subcellular dynamics in live cells, which is a critical advantage for cell biology and biomedical research^10-17^. However, current super-resolution techniques face significant trade-offs. Single-molecule localization microscopy (SMLM) requires ten thousand frames and is generally limited to fixed samples^13^. STED microscopy places stringent requirements on dye properties and laser power^18^, resulting in substantial phototoxicity for live-cell imaging. Therefore, techniques offering moderate resolution but faster imaging speeds and lower phototoxicity, such as structured illumination microscopy^19^ (SIM), super-resolution optical fluctuation imaging^14^ (SOFI), and super-resolution radial fluctuations^20^, are more compatible with live cell imaging.

While SIM is well-suited for live-cell imaging due to its low phototoxicity and fast acquisition^19^, its resolution is limited to roughly twice the diffraction limit and is often affected by reconstruction artifacts. To mitigate these issues, post-processing approaches such as deconvolution^21-23^ and deep learning^23-25^ have been developed to improve image quality. SOFI, on the other hand, offers a theoretically unlimited resolution increase by correlating higher-order fluorescence fluctuations and thus shows good momentum. However, for weakly fluctuating fluorescence signals or low signal-to-noise (SNR) images, hundreds of frames are still needed to suppress noise and reduce reconstruction artifacts^13,26,27^, and the image number demand for higher-order SOFI reconstruction may rise exponentially as well. Therefore, the application of self-blinking dyes^28^ and a two-step deconvolution method^29^ have been proposed to address the above issues separately that hinder the practicality of SOFI. Although recently SOFI has the potential to be used in applications comparable to SIM, robust imaging for higher-order SOFI remains a challenge^27^. Current super-resolution methods rely on sliding windows and multiple frames for real-time imaging^19,26,30^, which are sensitive to movement artifacts for live cells. To overcome these limitations, single-shot super-resolution techniques like instant structured illumination microscopy^31^ (iSIM) and image scanning microscopy^5^ (ISM) have been established. Although based on different imaging layouts, both depend on structured reconstruction and therefore suffer from noise and artifacts similar to SIM.

In this work, we present spatial polarization-induced fluorescence fluctuation imaging (SPIFFI), a single-shot super-resolution technique that exploits the polarization properties of fluorescent dipoles in their excited states. The spread function of a fluorescent dipole under polarized detection inherently exhibits a smaller waist size compared to the isotropic point spread function (PSF) typically modeled in conventional fluorescence microscopy. By introducing controlled polarization contrasts across multiple detection channels, SPIFFI utilizes physically constrained spatial fluctuations that enable instant resolution enhancement of at least 1.7-fold. These single-frame super-resolved images can be further processed through a dual-stage reconstruction strategy, achieving a final resolution of ∼80 nm without introducing detectable reconstruction artifacts. Beyond resolution improvement, the intrinsic polarimetry design of SPIFFI also enables direct readout of fluorescent dipole orientation and anisotropy. These added dimensions open powerful opportunities for investigating the spatiotemporal organization and functional anisotropy of biomolecules within their native cellular context.

## 2. Results

### 2.1 The working principle and verification of SPIFFI

Axial slices are acquired with 50 nm scanning steps. **(d)** The WF image of 160 nm nanorulers and the 3^rd^ order SPIFFI were reconstructed from a single WF frame. The intensity profile of the selected ruler yields an inter-marker distance of 163 nm. **(e)** Resolution evaluation using 100 nm diameter beads and 160 nm length nanorulers. The FWHM of lateral PSFs is calculated for beads and the distance between two marks is measured for nanorulers. For beads, lateral PSF size is 284 ± 8 nm (WF) and 166 ± 12 nm (SPIFFI). For nanorulers, the inter-marker distance is 156 ± 11 nm (SIM) and 163 ± 15 nm (SPIFFI). Distance values represent mean ± S.D.

Because most fluorescent dyes in biological imaging emit from dipolar excited states^32^, we exploit this property by splitting the emission into four detection channels, each detecting a distinct linear-polarization state (**Supplementary Section 1 - 3**). As illustrated in **Fig. 1a**, a single widefield (WF) image is decomposed into four polarized sub-images, each exhibiting spatial intensity variations due to the angular orientation of the fluorophore’s dipole. But we shift the fluctuations from the temporal to the spatial domain with the help of multichannel polarimetry, in contrast to conventional SOFI’s reliance on time-resolved fluctuation. However, to take advantage of SOFI in terms of noise suppression and contrast enhancement, we introduce background subtraction and virtual resampling on four-channel images for data augmentation, which is inspired by the Noise2Noise (N2N) and Noise2Fast (N2F) denoising models^33,34^ (**Supplementary Section 4**). First, background subtraction is based on the rolling-ball algorithm that removes uneven illumination and out-of-focus backgrounds. Next, because of the symmetrical shape of the point spread function (PSF) in the fluorescence image, the virtual resampling is performed by checkerboard down-sampling before recovering the image size by using the Sibson interpolation, creating a noise pair. In parallel, rotating the image by 90° and repeating the process yields another pair, increasing the receptive field of spatial neighbors. This step effectively expands the receptive field for neighboring pixels involved in the interpolation process, thereby enhancing the ability to capture subtle spatial fluctuations across polarization channels. Each channel thus produces five fluctuation images (original + 4 synthetic), resulting in 20 spatially fluctuating fluorescence images in total. Finally, we apply pre-deconvolution to improve the reconstruction quality and image shuffle to enhance the spatial fluctuations before performing autocorrelation computation for single-shot super-resolution reconstruction. In this way, a super-resolved temporal image is generated for each single frame without relying on time-lapse acquisition or sliding windows, thus avoiding motion artifacts commonly encountered in live-cell imaging. A detailed flowchart can be found in the **Supplementary Section 5**.

**Fig. 1.**
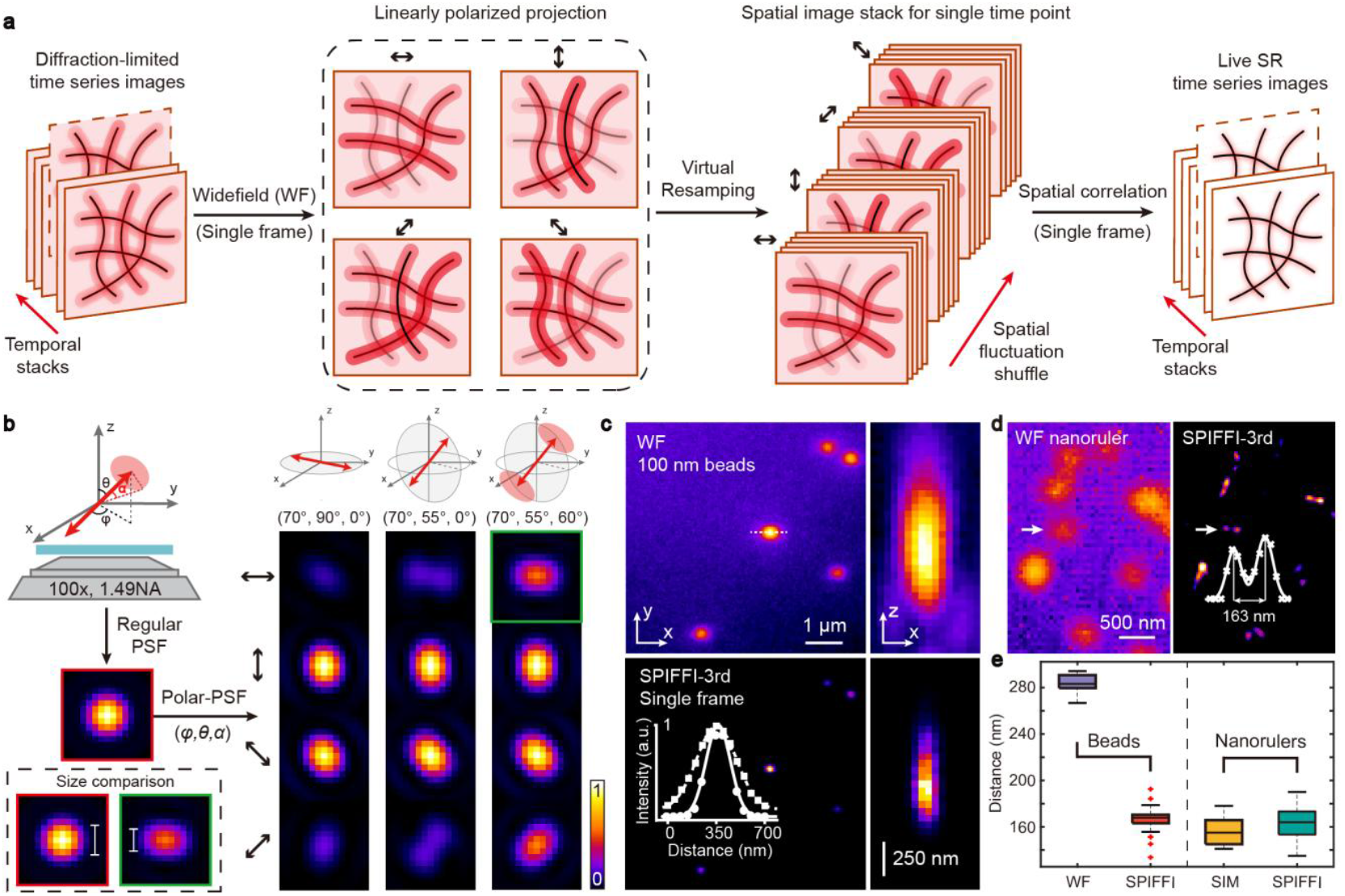
Working principle and resolution validation. **(a)** Workflow of SPIFFI based on spatial fluctuations captured across four polarization-resolved channels. **(b)** Simulated polarized spread functions of fluorescence dipoles with varying orientations (azimuthal angle *φ*, polar angle *θ*) and restricted wobbling (half-cone angle *α*). The slimmer waist of each polarized PSF indicates a smaller diffraction limit than the conventional isotropic PSF. The pixel size of each PSF image is 50 nm, the emission wavelength is 580 nm and all results are simulated under the same conditions. **(c)** Lateral and axial cross-sections of 100nm diameter beads (emission wavelength 580 nm). The upper panel shows a single WF image, and the bottom panel shows the corresponding SPIFFI reconstruction. The inset is a comparison of the intensity profile of the selected bead.

The resolution enhancement of SPIFFI arises from three aspects: (1) small size of polarized-PSFs; (2) polarization-modulated fluorescence fluctuations^35^ and (3) contrast enhancement benefiting from image denoising. As shown in **Fig. 1b**, The WF-PSF is usually approximated as a two-dimensional (2D) Gaussian shape for an isotropic point source with an emission wavelength of 580 nm, but for a dipole with three degrees of freedom (azimuthal angle *φ*, zenithal angle *θ*, and wobble angle *α*), polarized-PSFs are mostly elliptical and their waist size is naturally smaller than the WF-PSFs. This direct size reduction is also robust to the orientation and rotational mobility of the fluorescent dipole under the same simulation conditions, which directly ensures that the resolution of the polarized fluorescence image is smaller than the conventional diffraction limit (**Supplementary Section 6)**. When the dipole wobbles uniformly in a hard cone (i.e. restricted wobbling), which is common in fluorescent labeling, its rotational diffusion could also blur the polarized-PSF splitting caused by a small zenithal angle and reduce imaging ambiguity. In addition, the second and third points mentioned in the beginning usually work in parallel. SPIFFI ensures that the fluctuation analysis is performed directly on the same molecule, which shrinks the PSF directly and avoids image degradation caused by image drift. Moreover, virtual resampling and background subtraction make the background approximately follow a zero-mean distribution, so that the correlation of background noise is much smaller than that of the fluorescence signal, resulting in noise suppression and contrast improvement. This level of noise suppression is not achievable by methods such as Richardson–Lucy deconvolution, which iterates to noise-dominated artifacts^36^. Finally, the reduction of the PSF waist size is not isotropic, and it is strongly correlated with the dipole orientation, but the autocorrelation calculation is free from the trouble of channel selection and fully utilizes the spatial polarization-induced fluctuations.

We validated SPIFFI’s performance using 100 nm beads (emission wavelength 580 nm) and DNA nanorulers. **Fig. 1c** shows 3D fluorescence images of 100 nm diameter beads with an axial scanning step of 50 nm. The lateral full-width at half-maxima (FWHM) of the 3rd-order SPIFFI for the selected bead is 173 nm, which is about 1.7 times better than the size of the WF (294 nm). The axial resolution of SPIFFI is 385 nm, which is 1.8 times better than the wide-field 710 nm. For structural resolution, we imaged DNA nanorulers (SIM 160Y, 160 ± 5 nm in length) labeled with ATTO 565 at both sides in **Fig. 1d**, providing a precise two-point distance to benchmark the system’s spatial resolving capability. We successfully resolved the pattern of nanorulers and the distance of a selected nanoruler is 163 nm by double Gaussian fitting. But compared to the other methods, nanorulers cannot be resolved by commercial Airy-SCAN, and the result of SIM suffered from artifacts (**Supplementary Section 7)**. Importantly, SPIFFI reconstructions were obtained from single raw frames, preserving full temporal resolution for the image stack. The lateral FWHM of SPIFFI is 166 ± 12 nm for beads in **Fig. 1e**, while the FWHM of widefield (WF) is 284 ± 8 nm. The distance resolved by SPIFFI is 163 ± 15 nm for 20 pairs of nanorulers, and the distance resolved by SIM is 156 ± 11 nm. In addition, since fluorescent beads can be approximated as freely rotating dipoles, the polarization contrast among the four channels is minimal. Nevertheless, SPIFFI still achieves a 1.7-fold resolution improvement in this undesirable scenario. This suggests that for dipoles with restricted rotational mobility, the performance is expected to be no worse than this level (**Supplementary Section 8**). These results establish a solid proof-of-concept for SPIFFI, and we next demonstrate its practicality in biological experiments.

### 2.2 Single-shot super resolution imaging of fixed cells

SPIFFI is applied to fixed cells in this section to demonstrate the feasibility of biological imaging. We first perform 2D imaging on immunolabeled microtubules in fixed cos-7 cells (**Fig. 2a**). When WF images of each channel are averaged over 100 frames (**Fig. 2b**), the benefit of polarization measurements can be visualized easily. The intensity profile across two closed filaments is plotted by a solid curve and the contrast of the other structure is highlighted by a dashed box. SPIFFI reconstruction reveals fine structural details in the microtubule network and effectively suppresses out-of-focus background. At the same time, we also display the performance of several fluctuation-based super-resolution techniques (SACD^29^, SOFI-AC^6^, eSRRF^30^, and SOFI-XC^37^) and widefield image after Richardson–Lucy deconvolution (RL-deconv) on the same data set (**Fig. 2c**). Among these methods, both SACD and eSRRF produces a super-resolved image by only using 100 frames. However, benefiting from polarization measurements, SPIFFI shows similar resolution even on single-shot imaging. In contrast, eSRRF suffer from some structural imbalances because of the nonlinearity of fluorescent blinking and limited SNR of raw data. The SPIFFI enables instant and real-time super-resolution imaging without using any sliding window, which is at least two orders of magnitude faster than these techniques for showing temporal dynamics. To assess spatial resolution quantitatively, we fit intensity profiles between two closed filaments (red arrows) using a double-Gaussian model. SPIFFI resolves a 165 nm structure with a fitting R^2^ of 0.9959 (**Fig. 2d**), while RL-deconvolution fails to resolve it. Additionally, as a benchmark, eSRRF resolves similar structure (160 nm), and the same structure in SACD is around 171 nm but whose contrast is slightly weaker than both eSRRF and SPIFFI. Fourier ring correlation^38^ (FRC) analysis in **Fig. 2e** further confirms SPIFFI’s superior resolution: 166 nm by SPIFFI (1 frame), 153 nm by eSRRF (100 frames) and 161 nm by SACD (100 frames). Since both SPIFFI and SACD employ the same and best iterations of deconvolution for fairly comparation, it can be seen that polarization measurements contribute to more effective extraction of the target’s fine structure. Compared with pure RL-deconvolution with the same number of iterations, SPIFFI still maintains better structural resolution. SPIFFI also exhibits good reconstruction fidelity^39^ compared to other fluctuation-based techniques (Supplementary Section 9-10). Moreover, these FRC results align with the resolution calculated by decorrelation analysis^18^ in **Fig. 2c**, which is used for resolution assessment in the following sections because it can directly estimate the resolution of a single image instead of an image pair (**Supplementary Section 11)**.

**Fig. 2.**
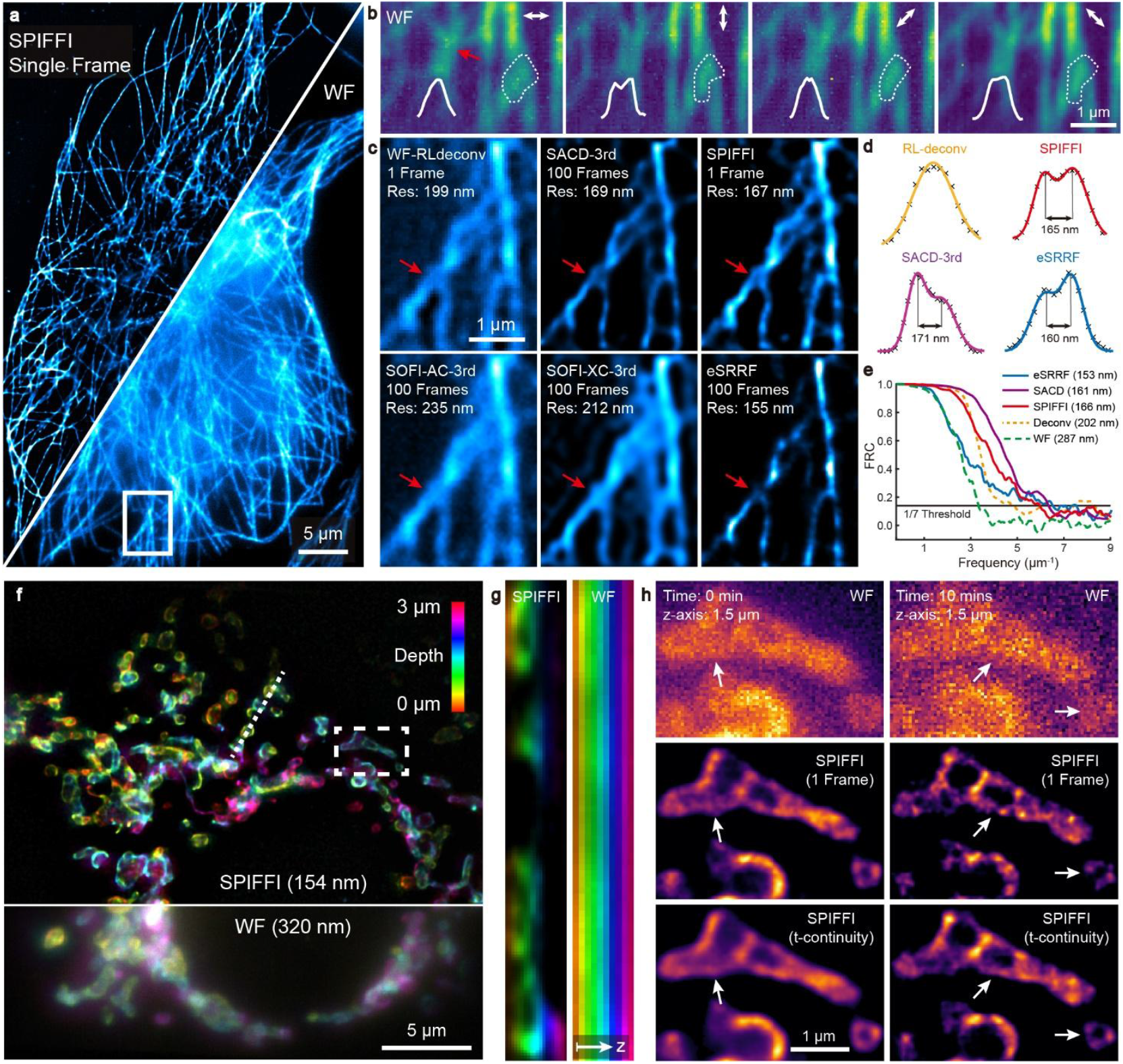
Single-shot super-resolution imaging of fixed cells. **(a)** 2D SPIFFI image of immunolabeled microtubules in a fixed COS-7 cell reconstructed from a single WF frame, shown alongside the corresponding WF image. **(b)** Averaged WF images (100 frames) from the four polarization channels. The red arrow highlights polarization-dependent intensity fluctuations arising from dipole orientation, and the second area shows the contrast difference presented by the polarization measurement. (c) Performance comparison of several fluctuation-based methods on the same regions. The red arrow marks a fine filament structure that is resolvable in the third-order SPIFFI image. The number of frames used in the reconstruction and resolution estimated by decorrelation analysis is marked in panels as well. **(d)** Lateral cross-section and double-Gaussian fitting for the structure marked by red arrows in (c). The minimum distance resolvable by the SPIFFI is 170 nm (R^2^ = 0.9917). **(e)** Image resolution is determined by Fourier ring correlation (FRC) using the 1/7 criterion. **(f)** 3D imaging of the mitochondrial outer membrane in a immunolabeled COS-7 cell. Ten axial planes (300 nm steps) are color-coded by depth. **(g)** Axial cross-section of the dashed region in (f). **(h)** SPIFFI reconstructions of the same region (white box) at high- and low-SNR conditions. Arrows indicate structures whose continuity is improved by a short window. The image fidelity of some arrow-marked structures is benefited from temporal continuity.

To demonstrate the advantages of SPIFFI in imaging speed and resolution enhancement, we extend its application to volumetric imaging of the outer mitochondrial membrane in fixed cos-7 cells (**Fig. 2f**). With the advantage in single-shot super-resolution, the intracellular mitochondrial network is visualized by ten axial scanning images with 300 nm axial steps and in color-coded depth. The improved 3D resolution helps us to successfully resolve the hollow structure and intertwined mitochondrial network in both lateral (**Fig. 2f**) and axial (**Fig. 2g**) planes, whereas WF images appear severely blurred. To improve image fidelity for low SNR images during longer-term imaging, we also implement a minimal sliding window strategy to fully utilize the temporal continuity of fluorescence images. As shown in **Fig. 2h**, single-shot SPIFFI resolves the outer mitochondrial membrane, but using a sliding window of three frames (±1 frame) could further enhance the integrity and smoothness of the outer membrane structure as white arrows marked. At the 10-minute timepoint, the membrane appears discontinuous in the single-frame SPIFFI result, but is restored with the short window strategy. SPIFFI acquires additional polarization dimension features without losing photons, making the utilization of photons more efficient in the super-resolution reconstruction.

### 2.3 Dual-stage reconstruction for further resolution improvement

In this section, we further improve resolution by performing a secondary reconstruction on the primary super-resolution images, combining both spatial and temporal fluctuations (**Fig. 3a**). All samples are labeled with blinking methods, because blinking enhances fluorescence fluctuations and thus reconstruction resolution; this also allows us to use SMLM data from the same samples as a benchmark. Each frame is first processed with third-order SPIFFI to obtain a primary super-resolution image, after which third-order autocorrelation SOFI is applied for a secondary reconstruction. For comparison, SMLM results are from the construction with ThunderSTORM^38^. This dual-stage approach robustly resolves 80 nm DNA nanorulers (Cy3B, GATTA-PAINT HiRes 80G) bearing three markers (**Fig. 3b**) by using only 1000 frames. But the resolution of the widefield image in **Fig. 3b** (SD, standard deviation) is only 254 nm. Intensity profiles of the same ruler in the dual-stage result and in SMLM yield similar inter-marker distances when fitted simultaneously with three Gaussian functions (**Fig. 3c**). The distances measured by SMLM and by the dual-stage reconstruction are 78.1 ± 4.2 nm and 78.9 ± 4.1 nm, respectively, which is in the range of the manufacturer’s specification (80 ± 5 nm; **Fig. 3d**). The dual-stage reconstruction achieves the equivalent of 9th-order SOFI while requiring only 1,000 frames, which is the number of frames normally used for second-order SOFI^13^. Moreover, it avoids the resolution degradation and cusp artifacts of high-order SOFI, because of separate low-order calculations from two different fluctuation sources and the direct PSF shrinkage of the polarimetry imaging **(Supplementary Section 13)**.

**Fig. 3.**
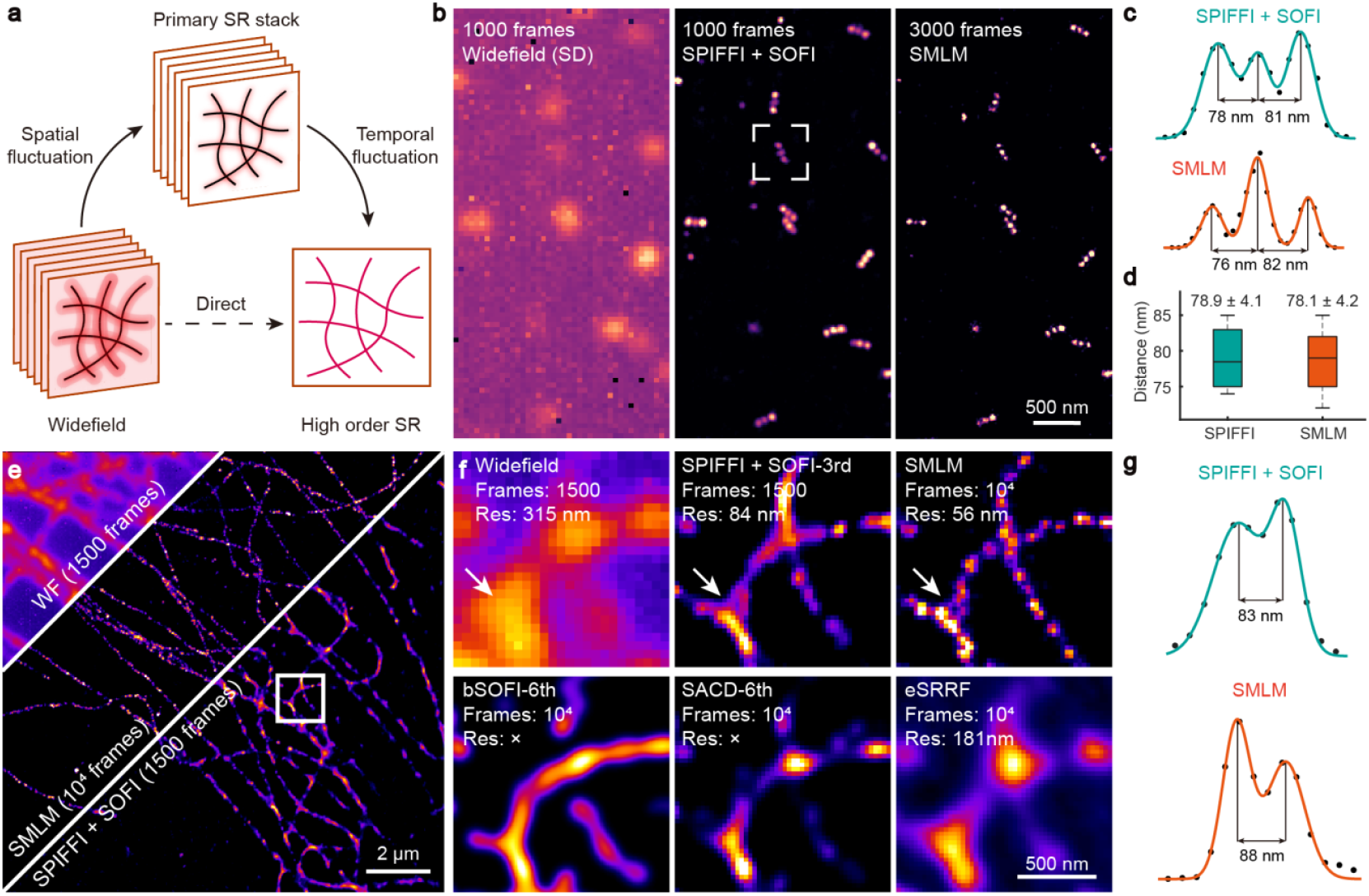
Dual-stage reconstruction achieves 80 nm resolution. **(a)** Workflow for the first (spatial fluctuation) and second (temporal fluctuation) reconstruction. **(b)** Comparison of widefield, dual-stage reconstruction and SMLM images of DNA nanorulers with 80 nm marker separation. The widefield image is the standard deviation of 1000 frames; both the two-stage construction and SMLM resolve all three labeled rulers. **(c)** The intensity profile of a representative nanoruler is fitted with three Gaussian functions simultaneously, confirming the 80 nm spacing. **(d)** Resolution evaluation using 80 nm spacing nanorulers. The distance between two markers measured by SMLM is 78.1 ± 4.2 nm and by two-stage construction is 78.9 ± 4.1 nm. **(e)** Validation of the dual-stage reconstruction on the fixed COS-7 microtubules immunolabeled with CF568. **(f)** Zoomed-in views of a selected area and comparison of different methods. The dual-stage reconstruction achieves approximately 80 nanometer resolution using only 1500 frames. **(g)** The intensity profile of a selected structure in (f). The dual-stage reconstruction shows similar distances as SMLM, which supports the image resolution from decorrelation estimation.

We will then validate the 80 nm resolution by imaging immunolabeled microtubules in a fixed COS-7 cell. The blinking and reversible switching of CF568 dye is induced by a STORM buffer^40^. Dual-stage reconstruction of 1,500 frames reproduces details comparable to SMLM reconstructed from 10,000 frames (**Fig. 3e**). Double-Gaussian fitting gives an 83 nm spacing between two microtubules, close to the 88 nm measured by SMLM. In addition, we want to emphasize the impact of image drift on long-sequence reconstructions, because a lateral drift of approximately 50 nm over 1000 frames (with an exposure time of 50 ms per frame in our case) is common in experiments. Therefore, we use a cross-correlation correction algorithm^41^ to compensate for the drift of the piezo stage. Considering the worst-case scenario under Nyquist sampling, the achievable resolution (i.e., <100 nm in our case) is fundamentally limited to no better than twice the drift distance. This prevents reliable recovery of 80 nm nano-rulers and the fine microtubule structure without drift correction (**Supplementary Section 14)**, even though the drift is smaller than the structures that we resolved.

### 2.4 Multidimensional super-resolution imaging for live cells

SPIFFI enables fast volumetric scanning, thereby avoiding motion artifacts from repeated exposures at the same depth in living cells **(Supplementary Section 15)**. As a demonstration, we image mitochondria in a live COS-7 cell labeled with MitoTracker-Red to highlight the advantages of fast 3D super-resolution imaging for resolving organelle interactions. Images are acquired at five axial planes, 250 nm apart, and rendered as a depth-coded maximum-intensity projection (MIP) in **Fig. 4a**. SPIFFI maintains high image quality over time in the first slowly moving region of interest (ROI-1) in **Fig. 4b** and proves resistant to photobleaching (**Fig. 4c, d**). SPIFFI also resolves the inner mitochondrial membrane that is not visible under WF imaging. Even when MitoTracker photobleaches with a time constant (τ^pb^) of 53 s, SPIFFI preserves a resolution of 178.4 ± 3.4 nm, whereas the corresponding WF resolution broadens to 479.0 ± 20.7 nm. In addition, mitochondrial dynamics (e.g. fission and fusion) are essential for cellular functional homeostasis^12,42,43^. SPIFFI resolves axial movements with 385.5 ± 13.7 nm resolution, showing that a local axial shift at the 30-s mark in ROI-2 could be misinterpreted as fission in 2D imaging alone (**Fig. 4e**). Similarly, mitochondrial fusion also occurs in 3D space. A tentative axial contact at different depths is observed in ROI-3 at 28 s (**Fig. 4f & g**). Such pre-fusion interactions serve as a quality control step, ensuring fusion with only healthy and compatible partners. After the fusion event at 35 s, the newly formed mitochondrion shifts axially to the same imaging depth. This indicates that 2D imaging alone can miss transient pre-fusion interactions occurring outside the focal plane and may misidentify the actual fusion timing.

**Fig. 4.**
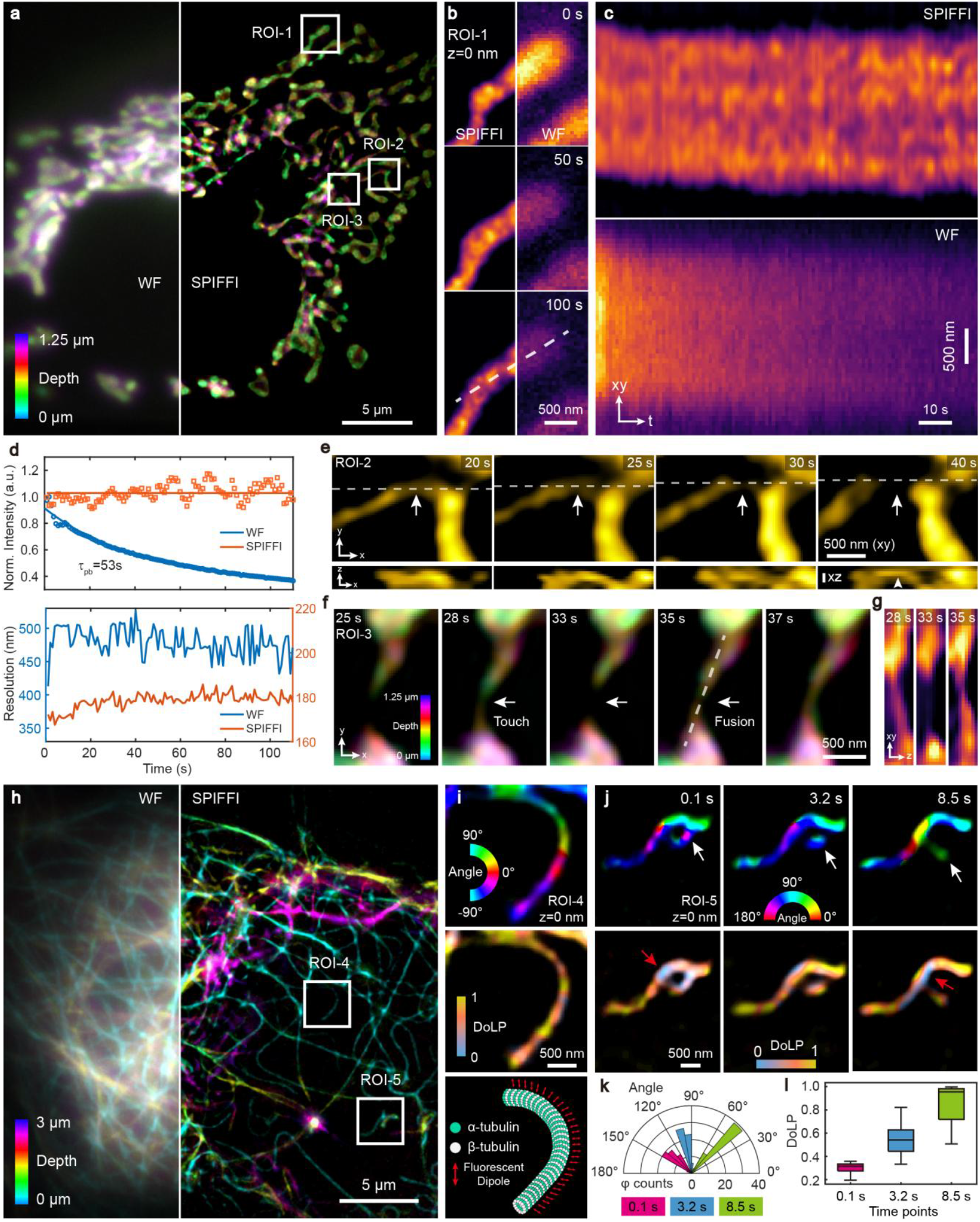
Single-shot and multidimensional super-resolution imaging for live cells. **(a)** Maximum intensity projection (MIP) of mitochondria in a live COS-7 cell labeled with MitoTracker Red. Axial depth of this cell is color coded. **(b)** Comparison of SPIFFI and WF in successive time points for first region of interest 1(ROI-1). **(c)** Space-time plots along the dashed line in (b). **(d)** The reconstruction quality of SPIFFI is verified from the image intensity (top) and resolution (bottom). **(e)** Time-lapse volumetric imaging of ROI-2 avoids the fission misinterpretation that arises in 2D imaging. **(f)** MIP of ROI-3 shows the dynamic mitochondrial fusion in 4D (xyz-t) imaging. The axial depth of this structure is color-coded. **(g)** Axial profiles of the ROI-3 structure in (f) at 28 s, 33 s, and 35 s reveal depth-dependent contact and fusion events. **(h)** 3D MIP of microtubules in a live COS-7 cell. The axial depth is color-coded by using the same colormap as in (a) and (f). **(i)** Fluorescent dipole parameters for the ROI-4 in (h). Top: orientation map of fluorescent dipoles; Middle: degree of linear polarization (DoLP); Bottom: organization of fluorescent dipoles along microtubules. **(j)** Time-lapse fluorescent dipole imaging of microtubules in ROI-5. White arrows in the orientation maps mark a dynamic structure of microtubules; red arrows in the DOLP images mark the anisotropy of probes. **(k)** Angular distribution and DoLP boxplots for the dynamic structures highlighted by white arrows in (j), quantifying orientation and anisotropy of fluorescent probes over time.

SPIFFI can not only achieve fast 4D (xyz-t) super-resolution imaging of living cells, but also reveal the orientation and anisotropy of fluorescent dipoles, offering critical insights into the molecular organization of biomolecules in the native cellular environment. Using a rhodamine-based fluorescent probe (Abberior LIVE 550) to label microtubules in live COS-7 cells, we collect ten axial planes with 300 nm steps (**Fig. 4h**). The MIP image reveals the microtubule network spanning the cell. Conjugation of the rhodamine dyes to tubulin-targeting ligands via short linkers leads to the formation of rigid complexes with minimal wobble angle, ensuring strong fluorescence polarization upon binding. Benefiting from the ordered orientation of fluorophores on the microtubule, the polarized channels capture fluorescence fluctuations with higher sensitivity, enhancing both resolution and contrast. Therefore, in the top panel of **Fig. 4i**, the orientation of fluorescent dipoles in the ROI-4 follows the curvature of the microtubule, remaining nearly perpendicular to filaments^44,45^. The degree of linear polarization (DoLP) map in the middle highlights the strong anisotropy of fluorescent dyes in the same region, which could be a combined effect of tilted dipoles, dense molecular packing, and molecular wobbling. Meanwhile, in the structure indicated by the white arrows in ROI-5 (**Fig. 4j**), the orientation of fluorescent dipoles changes as the microtubule filament transitions from a curved to a relative relaxed state, with measured angles of 136.6° ± 9.1°, 98.4° ± 5.6°, and 48.5° ± 5.5° at the respective stages in **Fig. 4k**. The corresponding DoLP rises accordingly in **Fig. 4l**, measured at 0.30 ± 0.04, 0.54 ± 0.12, and 0.85 ± 0.16, suggesting that the fluorescent dyes are tightly packed when the filament is curved, thereby reducing the anisotropy of the fluorescence signal. For dyes rigidly linked to targets, a decrease in DoLP typically occurs at filament nodes or bending regions, as exemplified by the structure marked with red arrows in the DoLP map of **Fig. 4j**. Supplementary experiments with FM4-64 in a giant unilamellar vesicle (GUV) and phalloidin-CF568 on actin filaments further verify SPIFFI’s dipole readout (**Supplementary Section 16**). FM4-64 inserts perpendicularly into the membrane on average, restricting wobble and producing relatively high DoLP, whereas dye aggregation on bundled actin reduces DoLP. By performing simultaneous 6D imaging (x, y, z, time, dipole angle, and DoLP) of various biological samples, we demonstrate broad potential for real-time studies of subcellular interactions, offering a powerful tool to probe dynamic molecular organization and functional anisotropy in live cells.

## 3. Discussion

The resolution limit in optical fluorescence microscopy is defined by the FWHM of the PSF, below which spatial details cannot be resolved. However, when fluorophores are modeled as fluorescent dipoles, polarization measurements reveal that the waist of their spread functions inherently has a smaller FWHM size, providing a physical foundation for spatial polarization-induced fluorescence fluctuation imaging (SPIFFI). SPIFFI achieves single-shot super-resolution imaging of live cells by directly reducing the PSF size through autocorrelation of spatially polarized fluctuations, enhancing SNR via background subtraction, and improving image fidelity with virtual resampling. Multidimensional dipole imaging simultaneously reveals the native molecular organization and anisotropy within cells. Furthermore, by integrating spatial and temporal fluctuations, SPIFFI achieves 80 nm resolution in fixed samples using only a comparable number of frames to conventional fluctuation-based methods, while effectively suppressing reconstruction artifacts.

Although SPIFFI has demonstrated excellent performance, several aspects still hold potential for further improvement. For instance, restricted fluorophore rotation (such as in immunofluorescence) and random labeling (such as with MitoTracker) do not significantly reduce the resolution of the reconstructed images, but they can affect the accuracy retrieval of dipole information. For optimal performance, we suggest using fluorescent probes with short, rigid linkers. Such probes minimize the rotational averaging of the fluorophore’s emission dipole, thereby preserving the intrinsic fluorescence anisotropy. This leads to a higher and more informative Degree of Linear Polarization (DoLP) in the widefield image, which enhances the contrast between the SPIFFI channels and improves the accuracy of the orientation measurement. Probes such as fluorescent proteins^44,46,47^ or phalloidin-dye conjugates^48,49^, which offer relatively rigid labeling, are therefore excellent candidates. In addition, the labeling density influences the quality of image reconstruction. Sparse labeling would lead to fragmented structures in super-resolved images, which is a general concern that affects the quality of all fluorescence imaging techniques. Membrane potential–based probes (e.g. MitoTracker) label both the inner and outer mitochondrial membranes simultaneously, which can also introduce cusp artifacts and compromise dipole orientation estimation under low SNR conditions. Therefore, to ensure reliable dipole reconstruction, it is advisable to label distinct structures with spectrally separated fluorescent probes. To address discontinuities of structures, novel deconvolution strategies can be applied to improve structural fidelity. Finally, the 3D imaging in this work is still performed by axial scanning with a piezo stage, so integration with directly volumetric imaging techniques, such as multiplane methods^50-52^, could significantly accelerate imaging speed and improve axial resolution. While this integration requires more sophisticated optical design and computational reconstruction, it is a feasible and promising future direction.

Overall, we provide a new insight into multi-channel polarimetry, which configuration is already widely used or easily implemented in many labs^48,49,53-55^, in the field of fluorescence super-resolution imaging. Through experiments on various samples (mitochondria, microtubules, actin filaments, nanorulers, and lipid membranes) and multiple resolution imaging modes (single-shot super-resolution, six-dimensional information, and dual-stage reconstruction), we demonstrate the flexibility and broad potential of SPIFFI in the biological application, especially in real-time and universal super-resolution imaging.

## 4. Methods

### 4.1 Imaging setup

The home-made polarization microscope was configured as follows: a 561 nm laser was converted into circularly polarized light with more than 95% ellipticity through a combination of a linear polarizer (LPVISE100-A, Thorlabs) and a quarter-wave plate (WPQ05M-561, Thorlabs). The sample was mounted on a sample holder attached to a nano-positioning system (Mad City Labs) to ensure precise position control. Fluorescence emitted from the sample was collected using a 100× oil immersion objective lens with a numerical aperture (NA) of 1.49 (UPLAPO100X, Olympus), which was optimized for polarization-sensitive imaging. Excitation light was reflected by a dichroic mirror (Di03-R561-t1, Semrock), and the emitted fluorescence was filtered using a combination of a 750 nm low-pass filter and a 561 nm notch filter to suppress residual excitation light. A variable waveplate (LCC1223-A, Thorlabs) was employed in the emission path to correct the polarization distortions introduced by the birefringence and differential transmission of the dichroic mirror coating. The fluorescence signal was divided into two optical paths by a non-polarizing 50:50 beam splitter (BS016, Thorlabs). In one path after the beam splitter, a half-wave plate (AHWP10M-580, Thorlabs) rotated the polarization state by 45°. Subsequently, both optical paths were further separated by polarizing beam splitters (PBS251, Thorlabs), yielding four distinct polarization channels in total. And all four channels were finally directed via mirrors onto the same scientific CMOS camera (Prime-95B, Photometrics) for image acquisition.

The commercial airy scan microscope was Zeiss LSM980 and the commercial SIM was Nikon N-SIMS. The LSM980 used a 63x, 1.40 NA oil immersion objective, and the objective lens for the SIM was an Apo TIRF 100x, 1.49 NA. The detector used in LSM980 was Axiocam 506 mono and in SIM was Photometrics Prime 95B. The wavelength of the laser used in the experiment depended on the fluorescent dye.

### 4.2 Image processing, analysis, and visualization

Super-resolution image reconstruction, data analysis and MIP visualization were all based on custom scripts in MATLAB. Visualization of SPIFFI results follows the linearization of bSOFI, but the visualization of wide-field images is unprocessed.The rolling ball algorithm, pseudo-color rendering, and 3D rendering in the supplementary video were based on the functions of FIJI^56^. The colormap of 3D rendering in Fig.2f was ‘hsv’, in Fig.4a, Fig.4f and Fig.4h are ‘biop-12colors’, and the DoLPs in Fig.4i and Fig.4j used the ‘isolum’^57^ colormap.

### 4.3 Cell culture

COS7 cells were cultured at 37 °C and 5% CO2 using Dulbecco’s modified eagle medium (DMEM) without phenol red (Gibco, Thermo Fisher Scientific), supplemented with 10% fetal bovine serum, 1% penicillin-streptomycin and 4 mM L-glutamine (all three from Gibco, Thermo Fisher Scientific). Cells were plated at a density of 50,000 per dish on 35 mm glass coverslips cleaned by plasma and coated with fibronectin.

### 4.4. Fixed-cell labeling

#### Microtubule and Mitochondrial labeling

COS-7 cells were pre-fixed in a pre-warmed buffer with 0.25 % glutaraldehyde (w/v) and 0.3 % (v/v) Triton X-100. After fixation with 4% paraformaldehyde (PFA) in PBS, cells were permeabilized with 0.25 % (v/v) Triton X-100 in PBS and blocked with buffer (2% (w/v) BSA, 0.2% gelatin, 10mM glycine, 50mM ammonium chloride NH_4_Cl in PBS pH 7.4) for 1 h. For microtubule experiments, cells were incubated overnight with monoclonal anti-α-Tubulin (T5168, Sigma, dilution 1:1000) at 4°C then washed three times for 5 mins each with blocking buffer. Cells were incubated with goat anti-mouse IgG (H+L) CF568 (SAB4600083, Sigma-Aldrich, dilution 1:1000) for 1 h at room temperature. For mitochondrial experiements, cells were incubated with rabbit anti-TOMM20 (ab186734, Abcam, 1:1000) overnight at 4°C and then incubated with goat anti-rabbit IgG AlexaFluor 568 (ab175471, Abcam, 1:1000) for 1 h at room temperature. The samples were then washed three times, 5 minutes each, with a blocking buffer.

#### Actin labeling

Cells were washed three times in PBS and fixed on ice with 4% paraformaldehyde in PBS for 15 minutes. After being washed three more times in PBS, the cells were permeabilized with 0.5% Triton X-100 in PBS at room temperature for 10 minutes, followed by three times PBS washes. The staining solution was prepared by diluting the stock solution (200 U/mL) 1:40 in PBS. Cells were incubated with Phalloidin-CF568 (00044, Biotium) for 20 minutes and washed three times with PBS.

### 4.5 Live-cell labeling

The MitoTracker Red CMXRos (M7512, Invitrogen) was diluted to a working concentration of 50 nM or 100 nM in DMEM supplemented with 5% fetal bovine serum and 1% penicillin–streptomycin. The cell culture medium was removed from the dishes, and a prewarmed staining solution was added. After an incubation period of 1 hour at 37 °C, the staining solution was replaced with fresh prewarmed medium, and the cells were imaged immediately.

The tubulin dye (Abberior Tubulin Live 550) was diluted to working concentrations of 50 nM in DMEM. The culture medium was removed, and a prewarmed staining solution was added together with 1 μM verapamil. After an incubation period of 1 hour at 37 °C, the staining solution was replaced with fresh prewarmed imaging medium, and the cells were imaged immediately.

### 4.5 GUV preparation

GUVs were generated by gel-assisted swelling by polyvinyl alcohol^58^ (PVA). A 5% PVA solution (w/w) was prepared in Milli-Q and heated to 90 °C in a water bath until fully dissolved. To construct an open growth chamber, a rubber O-ring was affixed to a cleaned coverslip using a silicone elastomer (Kwik-Cast, World Precision Instruments). Then 50 μL of the heated 5% PVA solution was coated on the coverslip and dried completely. Lipid mixtures (DPhPC, 850356, Avanti Research) dissolved in 10 μL chloroform (1 mg/mL) were deposited onto the dried coverslip and evaporated. Following lipid deposition, the chamber was filled with a 15 mM sucrose solution and prepared to match the osmolarity of the external imaging buffer. GUVs were allowed to form by spontaneous swelling at room temperature. For membrane visualization, 5 μM FM4-64 dye (T13320, Thermo Fisher Scientific) was dissolved in Milli-Q water.

## 5 Acknowledgments

A.R., W.G., and L.F. acknowledge funding from the European Research Council (grant 101020445— 2D-LIQUID) and were supported by the EPFL Center for Imaging through its 2022 Call for Interdisciplinary Projects in Imaging. We thank Dr. Arne Seitz and Dr. Thierry Laroche from the Bioimaging and Optics Platform (BIOP) at EPFL for helping with SIM and Airy Scan imaging. We thank Prof. Sylvie Roke and Mr. Zhi Li from the Laboratory for fundamental BioPhotonics (LBP) at EPFL for preparing GUV samples and critical reading of the manuscript.

## 6 Contributions

W.G. and A.R. initiated and designed the project; W.G. built the microscope, programmed the code, and analyzed the data; L.F. cultured and labeled cells; W.G. wrote the manuscript with the input from all authors; A.R. supervised the project; All authors discussed results and commented on the manuscript.

## 7. Author information

Laboratory of Nanoscale Biology, Institute of Bioengineering (IBI), School of Engineering (STI), École Polytechnique Fédérale de Lausanne (EPFL), Lausanne, Switzerland Wei Guo, Lely Feletti, Aleksandra Radenovic

